# vSPACE: Exploring Virtual Spatial Representation of Articular Chondrocytes at the Single-Cell Level

**DOI:** 10.1101/2024.02.07.577817

**Authors:** Chenyu Zhang, Honglin Wang, Seung-Hyun Hong, Merissa Olmer, Hannah Swahn, Martin K. Lotz, Peter Maye, David Rowe, Dong-Guk Shin

## Abstract

Single cell RNA sequencing technology has been dramatically changing how gene expression studies are performed. However, its use has been limited to identifying subtypes of cells by comparing cells’ gene expression levels in an unbiased manner to produce a 2D plot (e.g., UMAP/tSNE). We developed a new method of placing cells in 2D space. This system, called vSPACE, shows a virtual spatial representation of scRNAseq data obtained from human articular cartilage by emulating the concept of spatial transcriptomics technology, but virtually. This virtual 2D plot presentation of human articular cartage cells generates several zonal distribution patterns, in one or multiple genes at a time, reveling patterns that scientists can appreciate as imputed spatial distribution patterns along the zonal axis. The discovered patterns are explainable and remarkably consistent across all six healthy doners despite their respectively different clinical variables (age and sex), suggesting the confidence of the discovered patterns.

## I. Introduction

Single cell and single nuclear RNA sequencing (scRNAseq and snRNAseq) is fast becoming the transcriptional standard for assessing the molecular landscape of a tissue because it reveals the cellular heterogeneity of the tissue that is lost with bulk RNAseq. The huge datasets resulting from these studies (5-10K cells by 5-10K genes) present a major challenge for the graphical representation of the types of cells with a similar molecular signature that are imputed into functional properties. In tissues where there are well-defined molecular signatures, for example the immune system, clustering cells into distinct and biological subgroups using widely used Seurat software and UMAP visualization is distinctive and biologically meaningful. However, in tissues where cell types are not well defined at the molecular level because prior knowledge is not well developed, such as articular cartilage (AC), the outcome generates clusters of merging cell populations with nomenclature that is not meaningful to the chondrocyte biological community. Figure 1 illustrates the UMAP analysis published by our group using the Seurat program that identifies clusters with names of fibrocartilage, reparative, proliferative, homeostatic, and effector chondrocytes that do not align with the known properties of this tissue (Swahn et al. 2023). The common tSNE/UMAP approach often misidentifies functional clusters because unbiased selection of the most variable transcripts for the underlying Principal Component Analysis often fails to capture the most functionally important genes in part because the ontology specifically developed for selected subgroup of chondrocyte cells does not exist.

**Figure 1:**
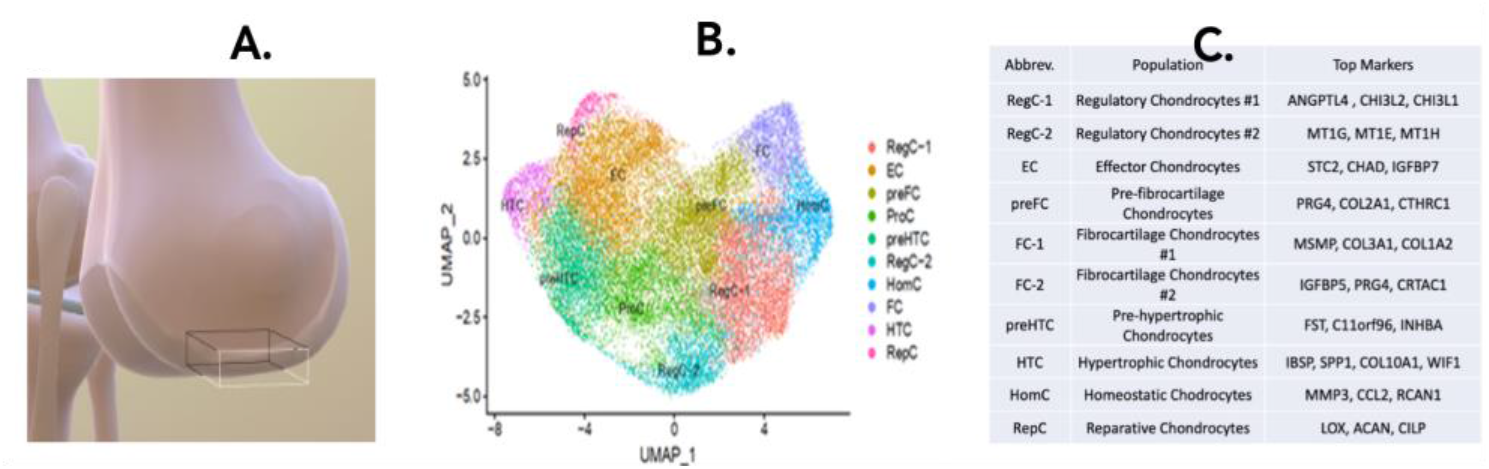
UMAP graphic (B) of the named clusters (C) obtained from a scRNAseq study (Swahn et al. 2023) of normal human femoral epiphyseal articular cartilage biopsy (A). The cluster names and key genes are inconsistent with the known biology of this tissue.

Here we report on a graphical approach vSPACE (<u>v</u>irtual <u>Sp</u>atial <u>A</u>rticular <u>C</u>artilage <u>E</u>xplorer) for representing an scRNAseq study of a tissue with well described histological organization, but which lacks the detailed cellular knowledge required for biologically meaningful clusters. It is designed to assist the AC community to develop meaningful clusters that eventually would be useful for spatial location of the clusters with the AC tissue. It is based on well-defined histological zone in the AC (superficial, transitional, middle, and deep, Figure 2A) that have been further refined at the transcriptional level by bulk RNAseq analysis of serial sections from the superficial to deep zone (Figure 2B).

**Figure 2:**
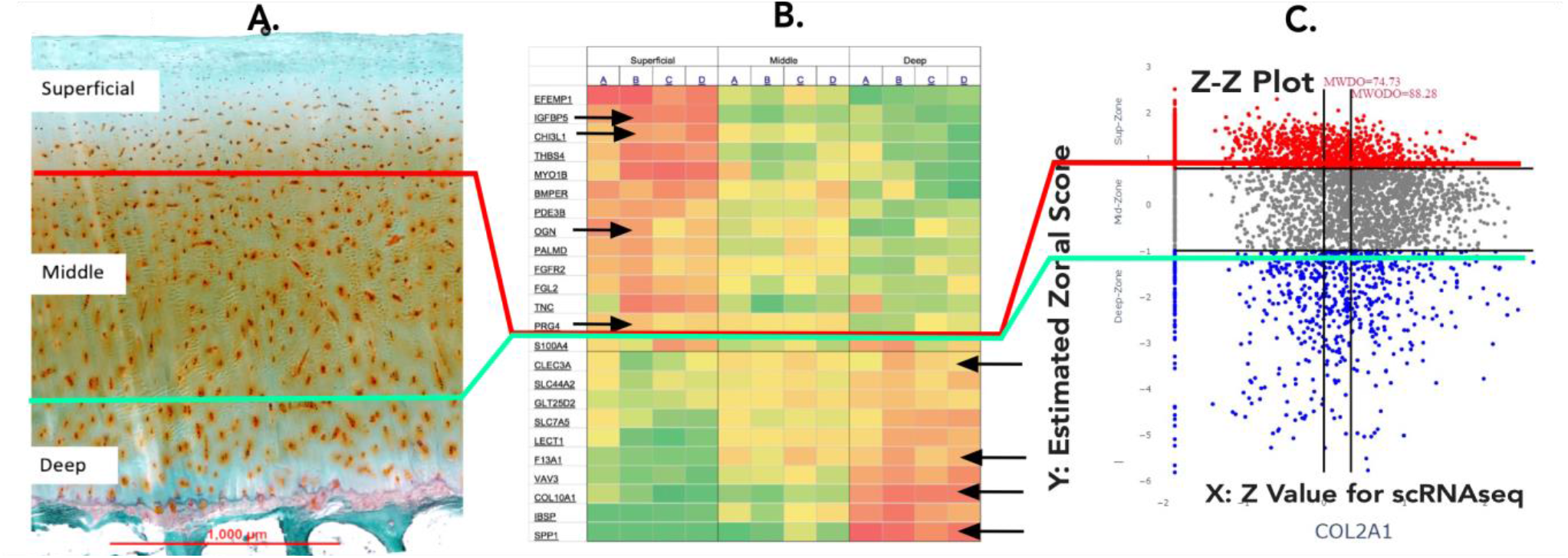
The histological zones of articular cartilage (A) that are mapped onto the heat map (Grogan et al. 2013) and used to position each cell onto the Z-Z spatial map (C). The arrows in B indicate the ZMG selected for the calculation that positions the cell on the Y axis of the Z-Z plot for the COL2A1 gene. The dots on the Y axis are cells that do not express Col2A1.

## II. Graphical Design Concept

Positioning each cell from scRNAseq study of AC within a 2D space (Figure 2C) is based on two determinants. The Y axis represents the estimate of the tissue’s top-to-bottom spatial positioning, meaning middle zone spreads out equidistance from Y’s value zero and Y’s higher positive value and lower negative value are mapped to, respectively, toward superficial and deeper zones. The Y value for each plotted cell reflects its similarity of gene expression to signal strength of the superficial or deep Zone Marker Genes (ZMG) that were selected from serial-section bulk RNA study as shown by the arrows in Figure 2B (Grogan et al. 2013). The calculation of Y for each cell in the study is derived from the difference of the weighted summed superficial ZMG score minus the weighted summed deep ZMG expression score. For the X axis the value represents the concerning gene expression strength in term of z-score, i.e. higher gene expression having large X axis value and X axis value 0 represents the gene’s population mean expression value and the X value being negative or positive means expression value being smaller or larger than the population average.

## III. Implementation

The developed vSPACE is a web-based toolbox for data processing, exploration, analysis and visualization of zonal gene expression patterns in articular cartilage (AC) cells. This web-based tool is written in the Python programming language (≥ 3.6.0) and takes advantage of the Dash (Dabbas 2021) for the pipelined and customizable visualization to produce publication-ready figures and tables.

### A. Input Data

The input data set needs to be pre-processed before being fed into the vSPACE portal. This step converts the raw data set into z-score format and computes all data matrices for a single cell including the averages of the gene expression values for each cell with the dropouts (genes that are not expressing), the averages of the gene expression value for each cell without the dropouts and the zonal labels. After this step, the processed data format also stores all the metadata, including the cell barcodes, gene symbol names, lists of marker gene sets, tissue types as well as the zonal names. Upon data input, a virtual spatial representation of the data points is then displayed in the interactive viewer at 2D space, also known as the “Z-Z plot” (Zone vs Z score). The following parts will illustrate how the Z-Z plot is constructed.

### B. Computing Zonal Score

Spatial markers for the AC cells are assumed to be known *a priori*, which we call the ZMG. How to find the most ideal ZMG for a single cell study performed on AC cells is an on-going research topic. In this study, we use the zonal markers produced by serial sections from the superficial to the deep zones analyzed with Mass Spec (Müller et al. 2014; PMID 25193283) and bulk RNAseq (Grogan et al. 2013; PMID 23124445) to compile a list of ZMG in the three major zones of the AC. Zonal Scores are calculated by Equation (1). We note that the explanation for this equation has been already detailed in our previous work (Zhang et al. 2022)

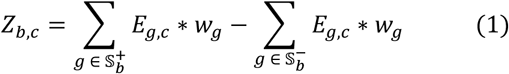

where *E*_*g,c*_ is the expression value of the gene *g* of the cell *c*, 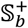 and 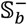 represent the positive regulation and negative regulation genes annotated for the biology defining the zonal index *b* respectively, and *w*_*g*_ is the weight factor of gene *g* calculated by Equation (2):

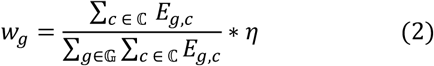

where *η* is a hyper-parameter determining the level of influence as a weight factor that can be chosen for each experiment. We typically set *η* = 0.2 as we have noticed from numerous empirical testings that this value tends to produce smaller *p-values* for BPs discovered. ℂ represents the set of all obtained cells from the data set and 𝔾 is the set of all available genes from the data set. All calculations are done before trimmed quantile normalization (Wang et al. 2022).

### C. Developing and Validating the Computational Spatial Map

For its two-dimensional placement of single cells, the Z-Z plot places each AC cell on the Y axis into three zones, superficial zone, middle zone and deep zone based the similarity of the expression profile to the serial bulk RNAseq where the z score for all cells expressing the gene determines the X axis value (Figure 2C). Note that cells not expressing the gene (dropouts) are placed on the Y axis line. The horizontal axis lines distinguish superficial zone and middle zone (upper horizontal line) and the middle zone and deep zone (lower horizontal line). The data points are then colored by the zonal scores: cells in superficial area (red dots), cells in middle area (grey dots) and cells in deep zone (blue dots). The vertical axis lines identify average value with dropouts (left vertical line) while the average value of expressing cells (dropouts excluded) is identified by the right vertical line. The average value of the data points that generated the two vertical lines are placed above the respective vertical lines (numbers in red). The gene symbol for the graphic is placed below the X-axis.

The program initially generates a Zonal Dot Count Table for the 9 Z-Z compartments (non-expressing, Z < 0, Z > 0 in the superficial, mid and deep zone). Subsequently it produces the Zonal Dot Percentage Table (Figure 3) that calculates percentage of all cells expressing the gene in each zone, excluding dropouts as well as the summed expression of the gene in all the cells in the scRNAseq study and the mean expression per total cell number and per expressing cells (non-expressing cells removed).

**Figure 3:**
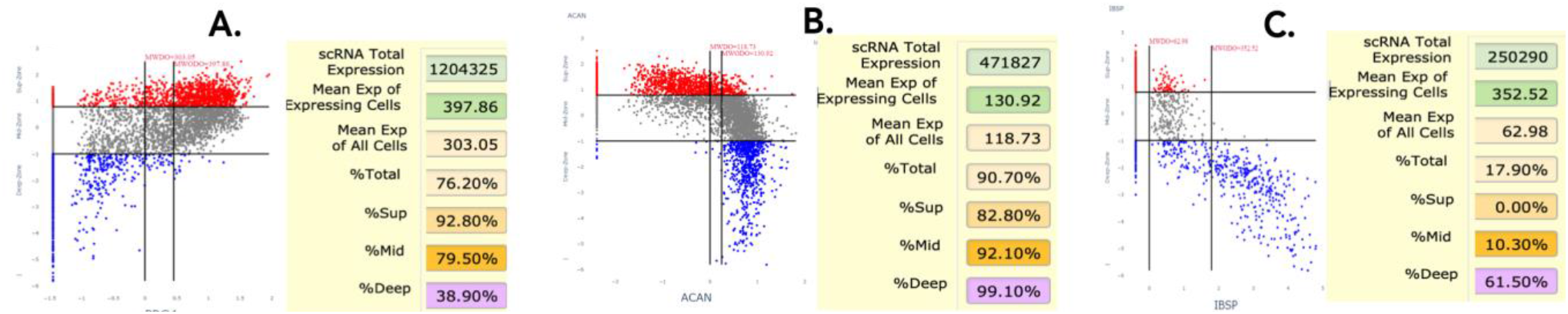
Z-Z plot of three genes whose distribution is well known by the AC community. A. Lubricin (PRG4) is produced by superficial zone cells. B. Aggrecan (ACAN) is a major matrix proteoglycan primarily located in the mid and deep zone. C. Integrin binding sialoprotein (IBSP) is major matrix protein of the deep zone. For each graph, the Zonal Dot Percentage Table shows mean level of total and per cell expression and percent cellular distribution within each histological zone.

Distinctive Z-Z patterns are produced when cells are selected for genes expressed in the three distinct zones. PRG4 produces the lubricin protein that is synthesized in the superficial zone and secreted into the joint fluid. The Z-Z pattern shows strong expression in the superficial and upper middle zone (Figure 3A). Aggrecan is a major proteoglycan that accumulates in the mid and deep zone which is reflected in the Z-Z pattern of strong expression in the two lower zones (Figure 3B). Integrin binding sialoprotein is a characteristic protein of the deep zone, and the Z-Z pattern matches this expectation by strong expression in the deep zone only (Figure 3C). Another validating observation is the Z-Z pattern of cell distribution of a single gene across multiple human donors selected by not having evidence of disease or trauma. Figure 4 illustrates the example of cells expressing the CLU gene across 5 human subjects (1 female, 4 male) ranging in age from 20 to 56. While the patterns are the same, the mean level and percent of expression can vary and could be an early marker of a subtle perturbation.

**Figure 4:**
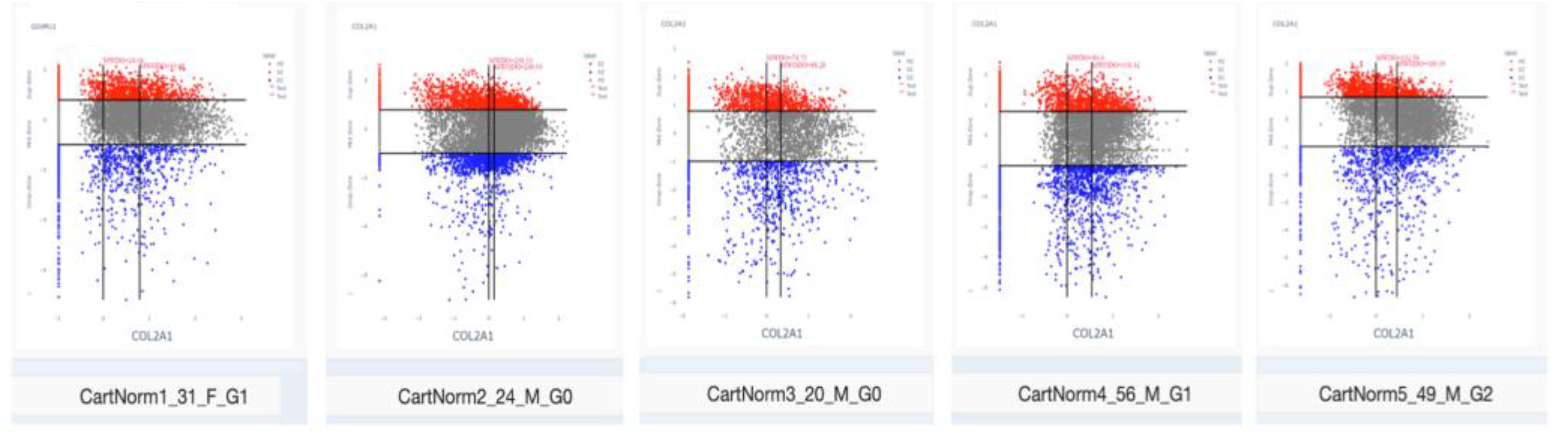
Consistency of the Z-Z pattern for COL2A1 (type II collagen) across AC samples obtained from healthy donors. Not shown in the comparison of the total and per cell expression across the donor set that does seem to vary by sex and donor age.

### D. Use of Spatial Mapping to Discover Clusters

Although the cellular histology of each zone has a similar morphology, vSPACE reveals a very heterogeneous population when viewed as molecular clusters. The analysis begins by selecting a well understood gene that is expressed in subpopulation of the AC such as PRG4, COL2A1, and IBSP (what we call “cluster sentinel genes”). Using the Spearman Coefficient feature of program, the genes most frequently expressed with the PRG4 sentinel gene are computed which in Figure 5 include CRTAC1 (cartilage acidic protein, an exported protein that like lubricin is secreted into the joint fluid). Figure 5A shows the Z-Z patten of PRG4 only (single gene mode) while Figure 5B overlays a heat map expression of CRTAC1 onto the PRG4 cells (in a co-expressing mode). Cells that strongly express both PRG4 and CRTAC1 have the bright orange color and are located in the superficial and top portion of the middle zone. Cells that are colored blue show low or no expression of both genes and are located in the mid percentage of cells strongly co-expressing the two genes so to deep zone. The Zonal Dot Count and Percent program then generates the number and that the percent enrichment within the cluster relative to the total cell population within each histological zone can be computed.

**Figure 5:**
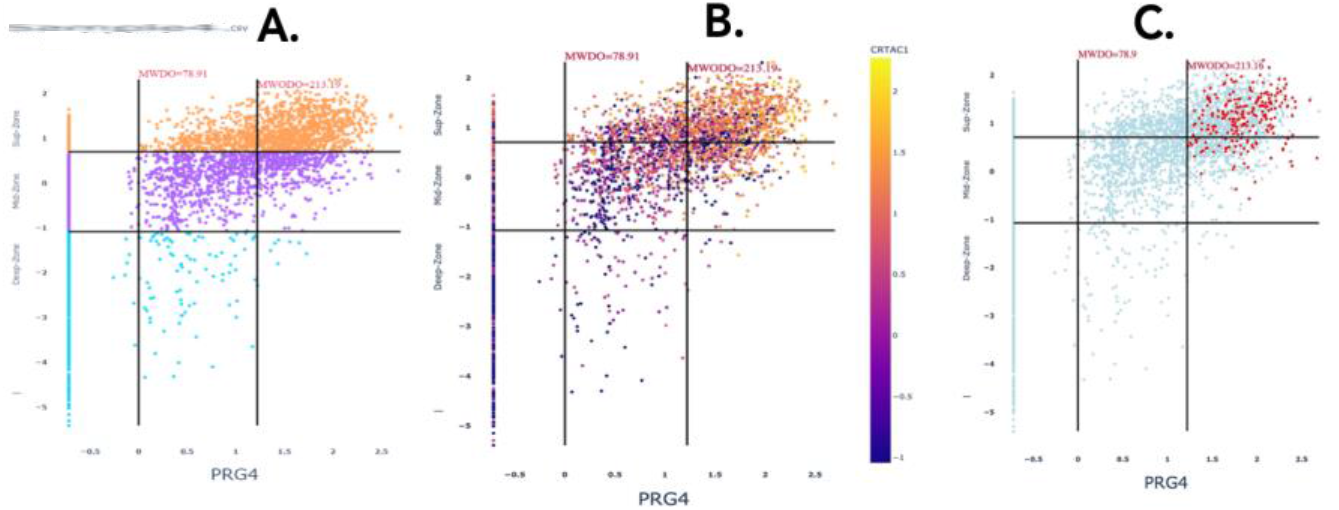
Graphical representation of cell clusters based on co-expression of marker genes. A. Z-Z plot of PRG4. B. Heat map of co-expressing CRTAC1 cells overlayed onto the PRG4 pattern showing high expression (Z >1) of both genes in the superficial zone. C. Overlay of cells with strong expressing (Z>1) of 3 genes (CRTAC1, DPT and ABI3BP) in PRG4 positive cells that show localization to the superficial zone.

Another feature uses a selection box (rubber banding) that encompasses the orange cells in the superficial zone that in turn generates an output file that tabulates the other genes that are expressed within the two-member cluster. From this list, other highly expressed gene(s) can be selected to further enrich for the specificity of the cluster. Figure 5C is designed to illustrate a multigene-co-expressed cluster, which in this case was PRG4-CRTAC1-DPT (dermatopontin, a secreted integrin binding protein) and ABI3BP (a secreted adaptor SH3 protein). The red dots within the entire PRG4 pattern are the cells that express the 4-member cluster. The user can perform another round of exploration real-time, e.g., Spearman correlation test or Mann-Whitney U test to find other genes enriched in the selected group from the web portal.

### E. Use by the AC biomedical community

The program is available to the public as a web-based tool at https://vspace.cse.uconn.edu. We envision this computation environment will have two major capabilities that will enhance the understanding of scRNAseq experiment of AC. The first is based by preloading an existing processed data files (normal or disease derived from different ages and sex) that are in the public domain and of excellent quality. These are designed to develop meaningful clusters and analytical pathways to supplant the current UMAP approach. The response rate of the program is dependent on the size of the data set (7071 cells x 445 genes vs 4450 genes) which in turn determines the complexity of the clusters that can be assessed with the tool. The second capability is to upload a study performed by the user which requires the conversion of the original data format to the file format the tool can ingest. This ingest file should be of the cvs format following the published column arrangement requirement which is available from the web tool. An example data ingest file is available in the web tool. Once in the program, the user has the option to input numerical values directly into the threshold fields or to alter the thresholds by dragging the horizontal axis lines to specify zonal markings. The Zonal distribution table updates automatically to reflect the thresholds set by the users. The user can enter one or more gene names to explore different gene pattern outcomes.

## IV. Conclusion

Our in-house use of the program environment has led us to appreciate the cellular complexity of a tissue that histologically appears to be very homogeneous. For example, a cluster based on strong Col2A1-COL3A1 co-expression reveals a cell population specializing in synthesis of multiple collagen types (COL5, COL6, COL9, COL11, COL12, COL15, COL16, COL27) while a COL2A-CTHRC1 cluster makes no COL transcripts but instead secretes matrix and growth factor modifying products. The Z-Z plot dramatically convey the difference between the pattern of COL2A1 between healthy and osteoarthritic cartilage with the percentage of cells expressing the transcript increasing from 60% to 100% (although a lower total number of cells) and the per cell expression increasing from 80 to 240. While bulk RNA sequencing does show a diminished COL2A1 mRNA, the scRNAseq provides a totally different perspective of how the AC chondrocyte responds to the disease environment. We predict that even more examples of early and late-stage disease will become evident when biological clusters are utilized for further interrogation. Finally, as marker gene combinations are discovered and made public, it will provide guidance for probe selection when designing and interpreting cell level spatial transcriptomic or proteomic studies. This product is designed to be a community resource that will be the foundation for a knowledge portal for the AC research community.

## Acknowledgment

Research reported in this work was supported in part by NIH Grant No. U54AR078664 and its NOSI supplement. Its contents are solely the responsibility of the authors and do not necessarily represent the official views of the NIH.

## References

Dabbas, Elias. 2021. Interactive Dashboards and Data Apps with Plotly and Dash: Harness the Power of a Fully Fledged Frontend Web Framework in Python–No JavaScript Required. Packt Publishing Ltd.

Grogan, Shawn P., Stuart F. Duffy, Chantal Pauli, James A. Koziol, Andrew I. Su, Darryl D. D’Lima, and Martin K. Lotz. 2013. “Zone-Specific Gene Expression Patterns in Articular Cartilage.” Arthritis & Rheumatism 65(2):418–28.

Müller, Catharina, Areej Khabut, Jayesh Dudhia, Finn P. Reinholt, Anders Aspberg, Dick Heinegård, and Patrik Önnerfjord. 2014. “Quantitative Proteomics at Different Depths in Human Articular Cartilage Reveals Unique Patterns of Protein Distribution.” Matrix Biology 40:34–45.

Swahn, Hannah, Kun Li, Tomas Duffy, Merissa Olmer, Darryl D D’Lima, Tony S. Mondala, Padmaja Natarajan, Steven R. Head, and Martin K. Lotz. 2023. “Senescent Cell Population with ZEB1 Transcription Factor as Its Main Regulator Promotes Osteoarthritis in Cartilage and Meniscus.” Annals of the Rheumatic Diseases 82(3):403–15.

Wang, Honglin, Pujan Joshi, Chenyu Zhang, Peter F. Maye, David W. Rowe, and Dong-Guk Shin. 2022. “RCom: A Route-Based Framework Inferring Cell Type Communication and Regulatory Network Using Single Cell Data.” Pp. 1–4 in Proceedings of the 13th ACM International Conference on Bioinformatics, Computational Biology and Health Informatics.

Zhang, Chenyu, Pujan Joshi, Honglin Wang, Seung-Hyun Hong, Riqiang Yan, and Dong-Guk Shin. 2022. “Pola Viz Reveals Microglia Polarization at Single Cell Level in Alzheimer’s Disease.” Pp. 1387–92 in 2022 IEEE International Conference on Bioinformatics and Biomedicine (BIBM).

